# Aging alters the role of basolateral amygdala in intertemporal choice

**DOI:** 10.1101/556894

**Authors:** Caesar M. Hernandez, Caitlin A. Orsini, Chase C. Labiste, Alexa-Rae Wheeler, Tyler W. Ten Eyck, Matthew M. Bruner, Todd J. Sahagian, Scott W. Harden, Charles J. Frazier, Barry Setlow, Jennifer L. Bizon

**Affiliations:** Department of Neuroscience, University of Florida, Gainesville, FL 32610, USA; Department of Psychiatry, University of Florida, Gainesville, FL 32610, USA; Department of Pharmacodynamics, University of Florida, Gainesville, FL 32610, USA

**Keywords:** aging, basolateral amygdala, optogenetics, decision making, delay discounting, impulsive choice

## Abstract

Aging is associated with an increased ability to delay gratification. Moreover, even when matched for performance, young and aged subjects recruit distinct brain circuitry to complete complex cognitive tasks. Experiments herein used an optogenetic approach to test whether altered recruitment of the basolateral amygdala (BLA), a brain region implicated in valuation of reward-cost contingencies, contributes to age-dependent changes in intertemporal decision making. BLA inactivation while rats deliberated prior to choices between options that yielded either small, immediate or large, delayed rewards rendered both young and aged rats less impulsive. In contrast, BLA inactivation after choices were made (during evaluation of choice outcomes) rendered young rats more impulsive but had no effect in aged rats. These data define multiple, temporally-discrete roles for BLA circuits in intertemporal decision making and implicate altered recruitment of BLA in the elevated preference for delayed rewards that is characteristic of advanced age.

**Impact Statement:** Basolateral amygdala (BLA) performs multiple, temporally-discrete functions during intertemporal decision making. Differential engagement of BLA contributes to the enhanced ability to delay gratification that is characteristic of advanced ages.

## Introduction

Intertemporal choice refers to decisions between rewards that differ with respect to both their magnitude and how far in the future they will arrive. Biases in intertemporal choice, whether manifesting as extreme impulsivity or patience, strongly associate with psychiatric disease. For example, greater preference for smaller, immediate rewards (greater impulsive choice) is a hallmark of attention deficit hyperactivity disorder and substance use disorders (Bickel et al., 2014; Hamilton et al., 2015; Patros et al., 2016; Crowley et al., 2017), whereas a pronounced preference for delayed gratification is characteristic of the eating disorder anorexia nervosa (Steinglass et al., 2012; Kaye et al., 2013; Decker et al., 2015). Independent of psychopathology, decision making associates with life outcomes and changes across the lifespan (Denburg et al., 2007; Boyle et al., 2012; Beas et al., 2015; Hess et al. 2015). Contrary to economic models predicting that older individuals should account for reduced time on the horizon in making intertemporal choices, healthy older adults actually exhibit a marked increase in preference for delayed outcomes (Green et al., 1996, 1999; Jimura et al., 2011; Löckenhoff et al., 2011; Mata et al., 2011; Samanez-Larkin et al., 2011; Eppinger et al., 2012). Although this pattern of choice behavior is sometimes characterized as “wisdom”, increased preference for delayed over immediate rewards may also be maladaptive. For example, biases toward delayed gratification in older adults could contribute to inappropriately conservative financial strategies that forgo expenditures necessary to maintain quality of life.

The neural circuits underlying age-associated changes in intertemporal choice remain poorly understood. Relevant to elucidating this circuitry is the fact that intertemporal choice is a multicomponent process that involves a series of temporally distinct cognitive operations (Rangel et al., 2008; Fobbs and Mizumori, 2017). Specifically, most decisions begin with representations of past choices, as well as some idea of the outcomes associated with each choice option. These representations are weighted by one’s motivation to obtain the choice outcomes at the time of the decision. A second phase of decision making occurs after a choice is made and involves evaluating the outcome to determine the degree to which it matches its predicted value. Feedback from this evaluative process can be used to adjust representations of the options to guide future choices. Both deliberation before a choice and outcome evaluation after a choice are supported by a network of brain structures that mediate reward processing, prospection, planning, prediction error, and value computations (Peters and Büchel, 2011; Orsini et al., 2015a; Bailey et al., 2016; Fobbs and Mizumori, 2017). The basolateral amygdala (BLA), which forms associations between cues or actions and their outcomes (Johansen et al., 2012; Wassum and Izquierdo, 2015), plays a central role in decision making and has been specifically implicated in both deliberation and outcome evaluation (Schoenbaum et al., 1998, 1999; Ghods-Sharifi et al., 2009; Peters and Büchel, 2011; Zuo et al., 2011; Grabenhorst et al., 2012; Zangemeister et al., 2016; Orsini et al., 2017). The BLA undergoes structural and functional alterations with advanced age, and BLA activity during intertemporal decision making is attenuated in aged rats (Lolova and Davidoff, 1991; Rubinow and Juraska, 2009; Rubinow et al., 2009; Roesch et al., 2012; Burke et al., 2014; Prager et al., 2016; Samson et al., 2017). It remains unclear, however, how age-associated changes in BLA neurobiology impact intertemporal choice.

Optogenetic tools have been employed previously to define temporally-specific roles of BLA during deliberation and outcome evaluation in young rats performing a decision-making task involving risk of punishment. Specifically, BLA inactivation at temporally discrete timepoints in the decision process shifted choice behavior in opposite directions (Orsini et al., 2017), highlighting multiple roles for BLA processing in risky decision making. The present study used a similar *in vivo* optogenetic approach to both define the role of BLA neural activity in intertemporal choice, and further to determine if the roles of BLA change across the lifespan.

## Materials and Methods

### Subjects

Young (6 months old, n=22) and aged (24 months old, n=18) male Fischer 344 x Brown Norway F1 hybrid (FBN) rats were obtained from the National Institute on Aging colony (Charles River Laboratories) and individually housed in the Association for Assessment and Accreditation of Laboratory Animal Care International-accredited vivarium facility in the McKnight Brain Institute building at the University of Florida in accordance with the rules and regulations of the University of Florida Institutional Animal Care and Use Committee and National Institutes of Health guidelines. The facility was maintained at a consistent temperature of 25° with a 12-hour light/dark cycle (lights on at 0600) and free access to food and water except as otherwise noted. Rats were acclimated in this facility and handled for at least one week prior to initiation of any procedures. Rat numbers reflect final group sizes after exclusion of misplaced injections. A subset of rats completed only some of the behavioral epochs due to lost headcaps and premature death, and some rats were excluded entirely for misplaced injections. Only the final numbers of rats included in each analysis is provided below.

### Surgical Procedures

Surgical procedures were performed as in our previous work (Orsini et al., 2017). Rats were anesthetized with isofluorane gas (1-5% in O_2_) and secured in a stereotaxic frame (David Kopf). An incision along the midline over the skull was made and the skin was retracted. Bilateral burr holes were drilled above the BLA and five additional burr holes were drilled to fit stainless steel anchoring screws. Bilateral guide cannulae (22-gauge, Plastics One) were implanted to target the BLA (anteroposterior (AP): −3.25 mm from bregma, mediolateral (ML): ±4.95 mm from bregma, dorsoventral (DV): −7.3 mm from the skull surface) and secured to the skull using dental cement. A total of 0.6 μL of a 3.5 × 10^12^ vg/ml titer solution (University of North Carolina Vector Core) containing AAV5 packaged with either halorhodopsin (CamKIIα-eNpHR3.0-mCherry, n=8 young and n=7 aged rats) or mCherry alone (CamKIIα-mCherry, n=4 young and n=4 aged rats) was delivered through the implanted cannulae at a rate of 0.6 μL per min per side. Stainless steel obdurators were placed into the cannulae to minimize the risk of infection. Immediately after surgery, rats received subcutaneous injections of buprenorphine (1 mg/kg) and meloxicam (2 mg/kg). Buprenorphine was also administered 24 hours post-operation, and meloxicam 48-72 hours post-operation. A topical ointment was applied as needed to facilitate wound healing. Sutures were removed after 10-14 days and rats recovered for at least 2 weeks before food restriction and behavioral testing began.

### In Vitro Electrophysiology

For *in vitro* electrophysiological verification of functional halorhodopsin (eNpHR3.0), young (n=2) and aged (n=2) rats underwent surgery as described above except that neither guide cannulae nor skull screws were implanted. Following a one month survival, rats were deeply anesthetized via i.p. injection of ketamine (75–100 mg/kg) and xylazine (5–10 mg/kg). The brain was rapidly cooled via transcardial perfusion with cold oxygenated sucrose-laden artificial cerebrospinal fluid (ACSF) containing (in mM): 205 sucrose, 10 dextrose, 1 MgSO_4_, 2 KCl, 1.25 NaH_2_PO_4_, 1 CaCl_2_, and 25 NaHCO_3_. Rats were then decapitated, brains extracted and coronal slices (300 μm) prepared using a Leica VT 1000s vibratome. Slices were incubated for 30 min at 37°C in oxygenated low-calcium ACSF containing (in mM): 124 NaCl, 10 dextrose, 3 MgSO_4_, 2.5 KCl, 1.23 NaH_2_PO_4_, 1 CaCl_2_, and 25 NaHCO_3_, after which they were transferred to room temperature for a minimum of 30 min prior to experimentation. During recording experiments, slices were bathed in ACSF containing (in mM): 125 NaCl, 11 dextrose, 1.5 MgSO_4_, 3 KCl, 1.2 NaH_2_PO_4_, 2.4 CaCl_2_, and 25 NaHCO_3_, maintained at 28-30°C. The pipette (internal) solution contained (in mM): 125 K-gluconate, 10 phosphocreatine, 1 MgCl_2_, 10 HEPES, 0.1 EGTA, 2 Na_2_ATP, 0.25 Na_3_GTP, and 5 biocytin, adjusted to pH 7.25 and 295 mOsm with KOH. BLA neurons were visualized using a combination of IR-DIC and epifluorescence microscopy using an Olympus BX51WI microscope and a TTL-controlled light source (X-Cite 110 LED light source, XF102-2 filter set, Omega Optical, excitation 540-580 nm, emission 615-695 nm, also used for *in vitro* activation of eNpHR3.0). Patch pipettes were prepared with a Flaming/Brown type pipette puller (Sutter Instrument, P-97) from 1.5 mm/0.8 mm borosilicate glass capillaries (Sutter Instruments) and pulled to an open tip resistance of 4–7 MΩ using internal solution and ACSF noted above. Electrophysiological recordings were performed using a Mutliclamp 700B amplifier and Digidata 1440A digitizer (Axon Instruments/Molecular Devices). Electrophysiological data were collected at 20 kHz and low-pass filtered at 2 kHz. Data analysis was performed using OriginLab and custom electrophysiology analysis software written by CJF.

Functionality of eNpHR3.0 was assessed in current-clamp configuration. Current was continuously delivered through the patch pipette to induce steady firing (1-10 Hz), and a 500 ms pulse of light was delivered through the objective lens to activate eNpHR3.0. At the conclusion of experiments, slices were transferred to 10% formalin (4°C, 24 hr) to allow for *post hoc* histological analysis. Slices were washed in PBS, permeabilized in PBS containing 0.1% Triton-X, and incubated in streptavidin conjugate with fluorophore (1:1000, 594 nm, ThermoFisher S32356). Slices were then mounted onto slides and coverslipped using VECTASHIELD. Morphological reconstruction was achieved by creating an all-in-focus maximum intensity projection of a Z-series acquired with a two-photon laser scanning epifluoresence microscope (810 nm excitation).

### Behavioral Testing Procedures

#### Apparatus

Testing was conducted in 4 identical standard rat behavioral test chambers (Coulbourn Instruments) with metal front and back walls, transparent Plexiglas side walls, and a floor composed of steel rods (0.4 cm in diameter) spaced 1.1 cm apart. Each test chamber was housed in a sound-attenuating cubicle and was equipped with a custom food pellet delivery trough fitted with a photobeam head entry detector (TAMIC Instruments) located 2 cm above the floor and extending 3 cm into the chamber in the center of the front wall. A nosepoke hole equipped with a 1.12 W lamp for illumination was located directly above the food trough. Food rewards consisted of 45-mg grain-based food pellets (PJAI; Test Diet, Richmond, IN, USA). Two retractable levers were positioned to the left and right of the food trough (11 cm above the floor). A 1.12 W house light was mounted near the top of the rear wall of the sound-attenuating cubicle. For optical activation of eNpHR3.0, laser light (561 nm, 8–10 mW output at the fiber tip, Shanghai Laser & Optics Century) was delivered through a patch cord (200 μm core, Thor Labs) to a rotary joint (1 × 2, 200 μm core, Doric Lenses) mounted above the operant chamber. At the rotary joint, the light source was split into 2 outputs. Tethers (200 μm core, 0.22 NA, Thor Labs) connected these outputs to the bilateral optic fibers (200 μm core, 0.22 NA, 8.3 mm in length; Precision Fiber Products) implanted in the BLA (Orsini et al., 2017). A computer running Graphic State 4.0 software (Coulbourn Instruments) was used to control the behavioral apparatus and laser delivery, and to collect data.

#### Behavioral shaping and initial training

The intertemporal choice task was based on a design by Evenden and Ryan (1996) and was used previously to demonstrate age-related alterations in decision making in both Fischer 344 (Simon et al., 2010) and FBN (Hernandez et al., 2017) rats (Figure 1). Rats were initially shaped to lever press to initiate delivery of a food pellet into the food trough and were then trained to nosepoke to initiate lever extension. Each nosepoke initiated extension of either the left or right lever (randomized across pairs of trials), a press on which yielded a single food pellet. After two consecutive days of reaching criterion performance (45 presses on each lever), rats began testing on the intertemporal choice task.

**Figure 1.**
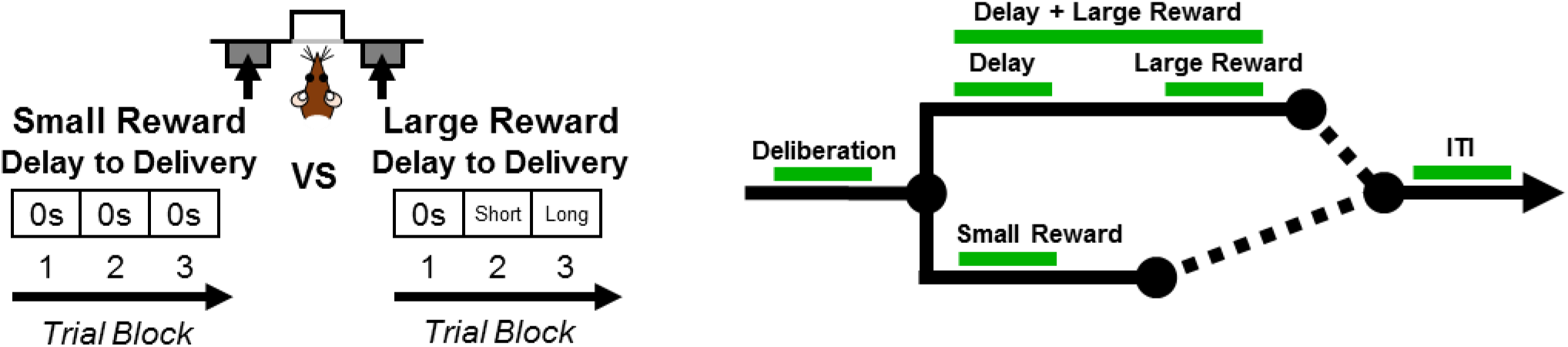
Schematics of intertemporal choice task and timing of light delivery. **A:** Schematic of the intertemporal choice task illustrating the choices and trial blocks across which the duration of the delay to the large reward increased. On each trial, rats were presented with two response levers that differed with respect to the magnitude and timing of associated reward delivery. Presses on one lever delivered a small (one food pellet), immediate reward, whereas presses on the other lever delivered a large (three food pellets), delayed reward. Trials were presented in a blocked design, such that the delay to the large reward increased across successive blocks of trials in a session. **B.** Schematic of a single trial in the intertemporal choice task, showing the task epochs during which light was delivered (represented by the green line). Using a within-subjects design, light was delivered during *deliberation* (when levers are presented until a choice is made); *small reward delivery; delay*; *large reward delivery*; *delay + large reward delivery*; and *intertrial interval (ITI*).

#### Intertemporal choice task

Each 60-min session consisted of 3 blocks of 20 trials each. The trials were 60 s in duration and began with a 10 s illumination of both the nosepoke port and house light. A nosepoke into the port during this time extinguished the nosepoke light and triggered lever extension. Any trials on which rats failed to nosepoke during this 10 s window were scored as omissions. Each 20-trial block began with 2 forced choice trials, in which either the right or left lever was extended, in order to remind rats of the delay contingencies in effect for that block. These forced choice trials were followed by 18 free choice trials, in which both levers were extended. For all trials, one lever (either left or right, counterbalanced across age groups) was always associated with immediate delivery of one food pellet (the small reward), and the other lever was associated with 3 food pellets (the large reward) delivered after a variable delay. Lever assignment (small or large reward) remained constant throughout testing. Within a session, the duration of the delay preceding large reward delivery increased across the three blocks of trials. The actual delay durations were adjusted individually for each rat on a daily basis, such that the percent choice of the large reward for each rat corresponded to approximately 100% in block 1, 66% in block 2, and 33% in block 3. On all trials, rats were given 10 s to press a lever, after which the levers were retracted, and food was delivered into the food through. If rats failed to press a lever within 10 s, the levers were retracted, lights were extinguished, and the trial was scored as an omission. An inter-trial interval (ITI) followed either food delivery or an omitted trial, after which the next trial began.

Rats were initially trained for 15 sessions on the intertemporal choice task. They were then lightly anesthetized and optic fibers (Thor Labs) were inserted into the guide cannulae such that they extended 1 mm past the end of the guide cannulae, and then were cemented in place. After recovery, rats resumed training but were now tethered to the rotary joint.

#### Effects of optogenetic inhibition during specific task epochs

The effects of temporally-discrete optogenetic inhibition of BLA were tested in both young and aged rats using a within-subjects design. Data from sessions occurring just prior to inactivation sessions (in which rats did not receive light delivery) were used as the baseline against which to compare the effects of BLA inhibition. Task epochs in which the BLA was inactivated included: *deliberation* [light delivery began 500 ms prior to illumination of the nosepoke light and continued until the rat pressed the lever (maximum of 10 s)]; *small reward delivery* [light delivery began when food was dispensed and remained on for 4 s]; *large reward delivery* [light delivery began when food was dispensed and remained on for 4 s], *delay* [light delivery began upon pressing the large reward lever and remained on throughout the delay (2-24s)]; *large reward delivery + delay* [light delivery began upon large reward choice and remained on until 4 s after the large reward was dispensed], *intertrial interval (ITI)*, [light delivery began 14 seconds after reward was dispensed and continued for 4 s]. Finally, the that the order in which the BLA was inactivated during different task epochs was randomized and counterbalanced across the two age groups.

### Vector Expression and Cannula Placement Histology

After completion of behavioral testing, rats were administered a lethal dose of Euthasol (sodium pentobarbital and phenytoin solution; Virbac, Fort Worth, TX, USA) and perfused transcardially with a 4°C solution of 0.1M phosphate buffered saline (PBS), followed by 4% (w/v) paraformaldehyde in 0.1M PBS. Brains were removed and post-fixed for 24 hours then transferred to a 20% (w/v) sucrose solution in 0.1M PBS for 72 hours (all chemicals purchased from Fisher Scientific, Hampton, NH, USA). Brains were sectioned at 35 μm using a cryostat maintained at −20°C. Sections were rinsed in 0.1M TBS and incubated in blocking solution consisting of 3% normal goat serum, 0.3% Triton-X-100 in 0.1M TBS for 1 hour at room temperature. Sections were then incubated with rabbit anti-mCherry antibody (ab167453) diluted in blocking solution at a dilution of 1:1000 (72 hours, 4°C). Following primary incubation, sections were rinsed in 0.1M TBS and incubated in blocking solution containing the secondary antibody (donkey anti-rabbit conjugated to AlexaFluor-488, 1:300) for 2 hours at room temperature. After rinsing in 0.1M TBS, sections were mounted on electrostatic glass slides and coverslipped using Prolong Gold containing DAPI (Thermo Fisher Scientific, Waltham, MA, USA). Slides were sealed with clear nail polish and sections were visualized at 20X using an Axio Imager 2 microscopy system (Carl Zeiss Microscopy, LLC, Thornwood, NY, USA) to assess mCherry expression in BLA neurons. Cannula placements and mCherry expression were mapped using a rat brain atlas (Paxinos and Watson, 2005). Decisions regarding inclusion/exclusion of rats based on cannula placements and mCherry expression within the BLA were conducted by an observer for whom rats’ behavioral performance was masked.

### Experimental Design and Statistical Analysis

#### Evaluation of age differences in intertemporal choice under baseline conditions

Raw data files were extracted using a Graphic State 4.0 analysis template that was custom-designed to extract the number of lever presses on each lever (large or small rewards) during forced and free choice trials in each trial block. First, age differences in intertemporal choice performance were tested by analyzing the actual delays used to achieve the target 100%, 66% and 33% choice of the large reward in blocks 1, 2 and 3, respectively. Actual delays were compared using a mixed-factor ANOVA, with age (2 levels: young and aged) as the between-subjects factor and block (3 levels: blocks 1-3) as the within-subjects factor. Second, the *choice indifference point*, or the delay at which a rat showed equivalent choice of the small and large reward, was calculated and compared between young and aged rats. Choice indifference points were calculated by fitting a trendline to each rat’s percent choice of the large reward at each delay block. The slope-intercept formula, y=mx+b (where “y” is percent choice or the large reward, and “x” is delay), was then used to solve for the number of seconds (x) at which y=50% choice of the large reward (the delay at which the rat was equally likely to choose the large or small reward). Choice indifference points were compared between young and aged rats using an independent samples t-test. For all statistical analyses and reported results, alpha was set to 0.05, η^2^ and Cohen’s *d* were used to report the effect size for mixed-factor ANOVAs and t-tests, respectively, and 1-β was used to report the observed power.

#### Evaluation of BLA inactivation on intertemporal choice

Power analyses based on data from an initial cohort of rats (n=3) were used to determine sample sizes necessary to evaluate the effects of BLA inactivation on choice behavior. These analyses indicated the presence of large effect sizes (greater than 1.0), and that n=6 rats should be sufficient to detect effects of BLA inactivation, with a power to detect significant differences of 0.95. The effects of light delivery were tested separately for each task epoch (*deliberation, small reward delivery, large reward delivery, delay, delay + large reward delivery*, and ITI). For each of these task epochs, comparisons were made using a mixed factor ANOVA (age × delay block × inactivation condition), with age as the between-subjects factor (2 levels: young and aged), and delay block (3 levels: delay blocks 1-3) and inactivation condition (2 levels: laser on or off) as within-subjects factors. To better understand significant main effects or interactions, *post hoc* analyses were conducted in each age group separately using a repeated-measures ANOVA (block × inactivation condition). Note that for those epochs in which effects of BLA inhibition during the delay were tested, data analyses were confined to blocks 2 and 3 as no delay was used in block 1.

#### Evaluation of choice strategy resulting from BLA inactivation

Additional analyses were conducted to better elucidate the shifts in young rat choice performance following BLA inactivation during deliberation and small reward outcome. Graphic State 4.0 templates were created to assess trial-by-trial choices during baseline and BLA inactivation sessions for the deliberation and small reward epochs. Trials were categorized based on choices made on the previous trial. For the deliberation epoch, trials were categorized as “small-<u>shift</u>-to-large” or “large-<u>stay</u>-on-large”. For the small reward delivery epoch, trials were categorized as “large-<u>shift</u>-to-small” or “small-<u>stay</u>-on-small”. The number of trials in each category was divided by the total number of trials in that session and expressed as a percentage. For each task epoch, percentages were compared using paired-samples t-tests comparing baseline and inactivation condition.

#### Effects on other task performance measures resulting from BLA inactivation

Other task measures were compared between BLA inactivation and baseline conditions in task epochs in which BLA inactivation produced significant changes in choice behavior. Specifically, on free choice trials, response latency (the time between lever extension and a lever press) was compared. Previous work shows that response latencies can differ for large and small reward levers (Hernandez et al., 2017) and hence analyses were conducted separately for each lever using data from delay block 2, during which rats made roughly equivalent numbers of choices on each reward lever. Response latency and total number of trials completed were compared using a mixed factor ANOVA (age × inactivation condition).

## Results

### Electrophysiological confirmation of light-induced inhibition of BLA neurons expressing eNpHR3.0

Virally-transduced (mCherry-positive) neurons in the BLA were visualized and targeted for whole cell patch clamp recordings using a combination of epifluorescence and differential interference contrast microscopy (Figures 2A-C). Consistent with previously published work that employed identical methodology (Orsini et al., 2017), mCherry-positive neurons (n=9 young and n=7 aged) were robustly hyperpolarized in response to 500 ms of light, and this hyperpolarization reliably eliminated action potential firing in cells (Figure 2D). *Post-hoc* morphological characterization confirmed that light-sensitive BLA neurons exhibited morphological characteristics consistent with pyramidal cells (i.e., large soma with numerous spiny dendritic arborizations, Figure 2C).

**Figure 2.**
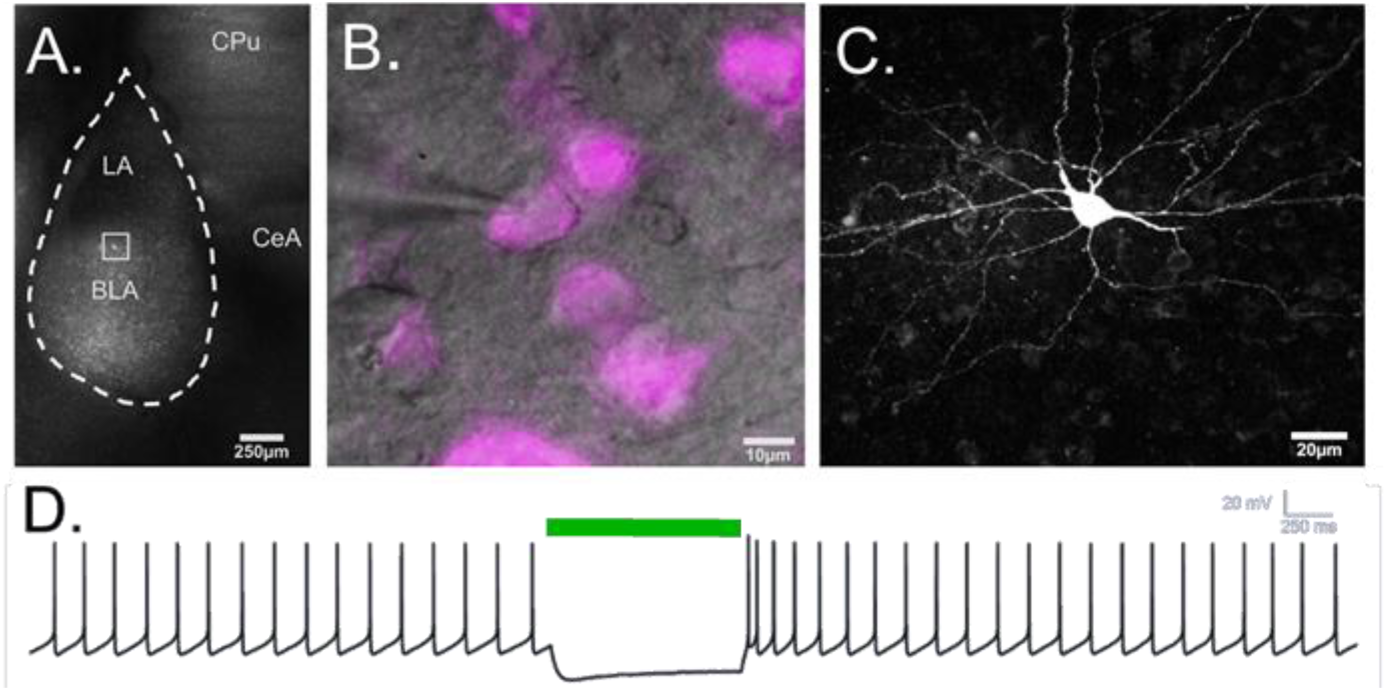
Functional inhibition of BLA pyramidal neurons via activation of halorhodposin in aged tissue. **A**. CaMKIIα-driven eNpHR3.0/mCherry was virally delivered into the BLA of young and aged rats. **B**. CamKIIα neurons were targeted for study by their fluorescence using a combination of DIC (gray) and epifluoresence (magenta) microscopy. **C**. A two-photon reconstruction of a biocytin-filled eNpHR3.0-expressing BLA neuron demonstrates multiple primary dendritic branches and spiny dendritic arborizations typical of BLA pyramidal neurons. **D**. Representative current-clamp trace from an aged BLA neuron demonstrates that a brief pulse of green light reversibly hyperpolarizes the neuron to silence firing.

### Fiber placement and AAV transduction

Expression of mCherry was used to confirm viral transduction in the BLA of rats used in behavioral studies that were injected with either AAV5-CamKIIα-eNpHR3.0-mCherry (AAV-eNpHR3.0, black circles in Figure 3) or AA5-CamKIIα-mCherry alone (AAV-control, white circles in Figure 3). Cannula placements were centered in the BLA, and the brain volumes transduced by AAV-eNpHR3.0 and AAV-control (calculated from the atlas of Paxinos & Watson, 2005) were comparable in young and aged rats.

**Figure 3.**
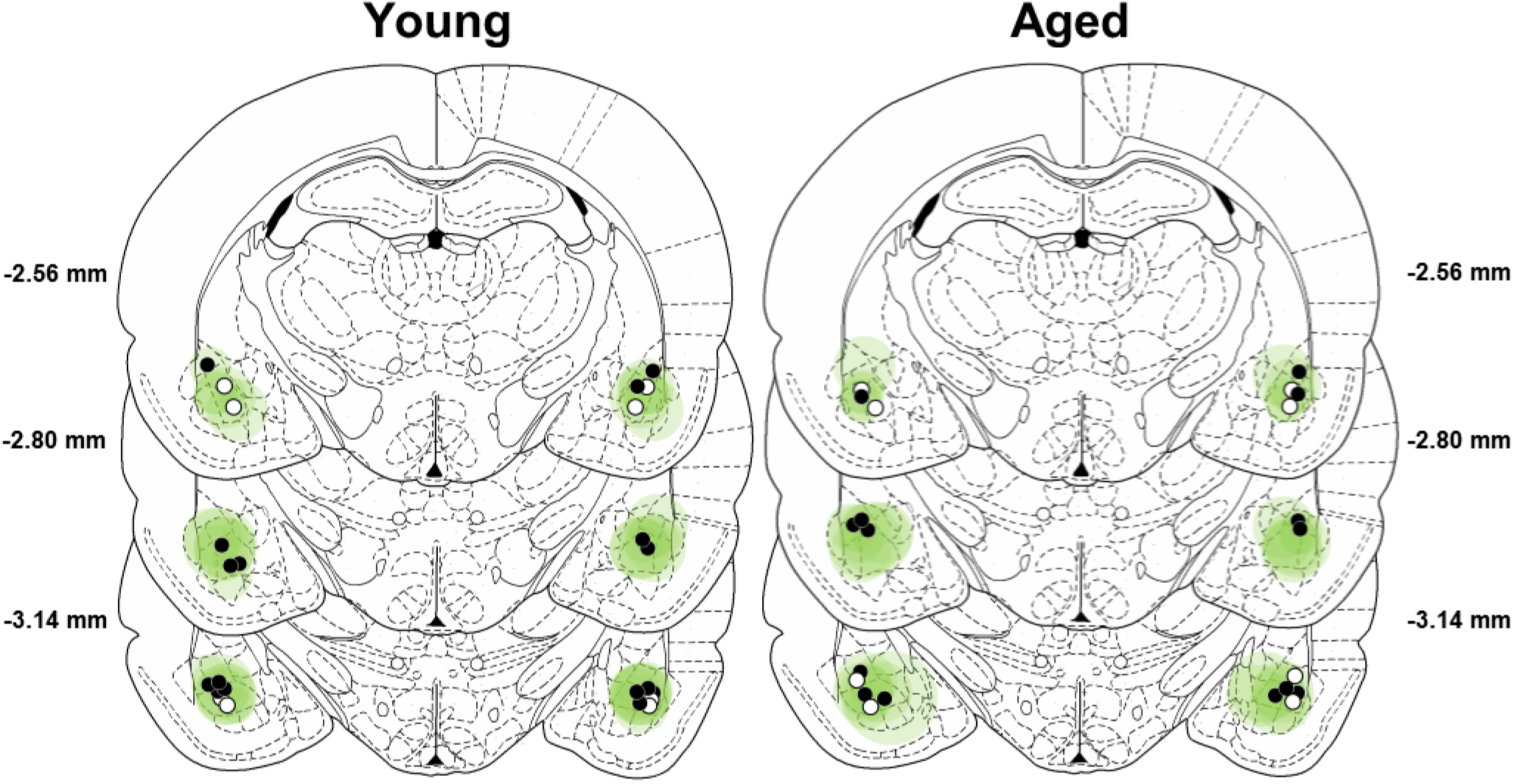
Verification of viral expression and fiber optic placements. The extent of viral expression in young (left) and aged (right) rats is depicted in green. Darker green indicates areas of greater viral expression (epicenter of the BLA). Lighter green indicates less viral expression (margins of the BLA). Filled black circles represent optic fiber placements in the experimental groups, and open black circles represent optic fiber placements in the control groups. Viral expression and fiber placements are mapped to standardized coronal sections corresponding to −2.12 mm through −3.30 mm from bregma according to the atlas of Paxinos and Watson (2005).

### Effect of age on intertemporal choice performance

Previous work shows that aged rats display attenuated discounting of delayed rewards (Simon et al., 2010; Hernandez et al., 2017). Therefore, prior to inactivation sessions, delays were adjusted on an individual basis to ensure that all rats’ choice performance was in the same parametric space (Figure 4A). This approach allowed a full and comparable range of effects from BLA inactivation to be observed in both young and aged rats, without concern for ceiling or floor effects. Figure 4B shows the actual delays used in the second and third blocks to achieve roughly 66% and 33% choice of the large reward, respectively, plotted as a function of age. A two-factor ANOVA (age × block) used to compare the actual delays indicated the expected main effect of block (F_(2,26)_ = 18.685, p<0.001, η^2^=0.606, 1-β=0.930), as well as a main effect of age (F_(1,13)_ = 6.402, p=0.025, η^2^=0.330, 1-β=0.648) and an age × block interaction (F_(2,26)_ =6.913; p=0.004, η^2^=0.347, 1-β=0.891). *Post hoc* analyses comparing the actual delays of young and aged rats in blocks 2 and 3 indicated that aged rats required longer delays than young to achieve comparable preference for large vs. small rewards (Block 2: t_(13)_=−2.234, p=0.044, Cohen’s *d*=1.114, 1-β=0.480; Block 3: t_(13)_=−2.660, p=0.020, Cohen’s *d*=1.328, 1-β=0.625). Consistent with this analysis, aged rats in comparison to young rats showed a greater indifference point (the delay at which rats showed equivalent preference for large and small rewards; t_(13)_ = −2.168, p=0.049, Cohen’s *d*=1.080, 1-β=0.457; Figure 4C).

**Figure 4.**
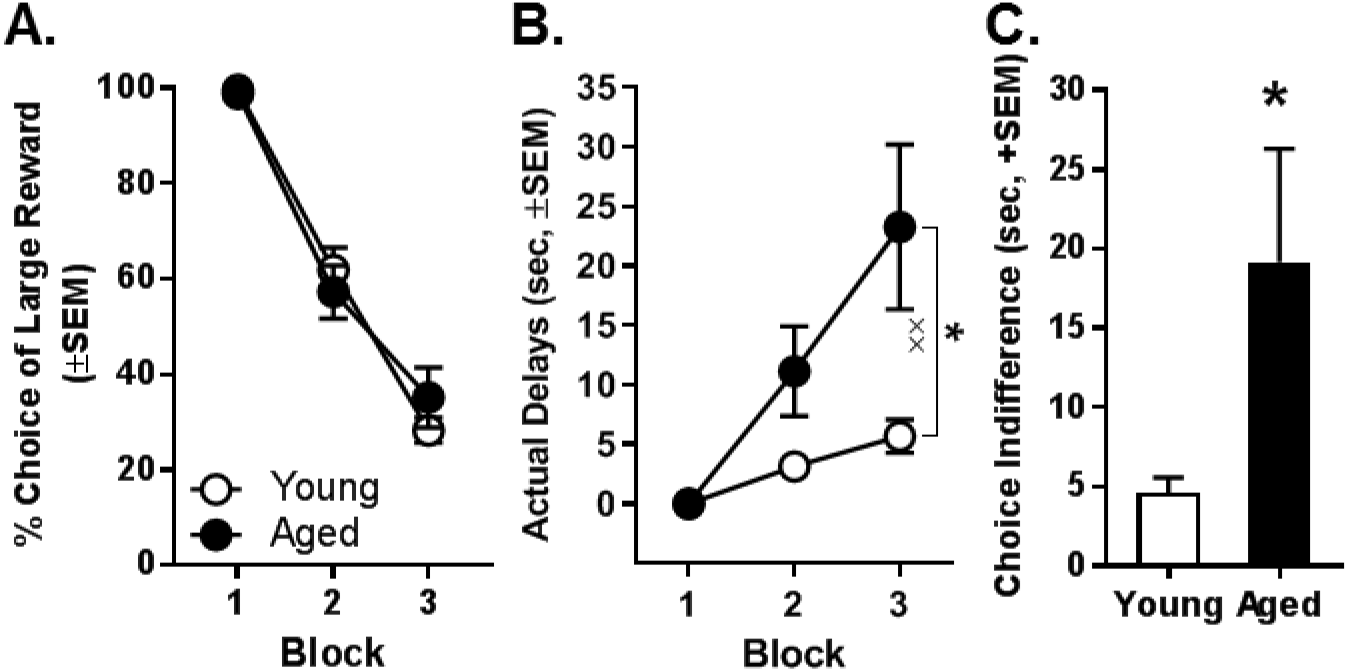
**A:** Mean percent choice of the large reward in young and aged rats prior to initiation of any BLA inactivation experiments. Note that delays to large reward delivery were adjusted individually for young (n=8, open circles) and aged (n=7, closed circles) rats in order to place all rats in the same parametric space. **B:** Mean actual delays required to achieve the comparable young and aged rat choice performance shown in panel A. Aged rats required longer delays in Blocks 2 and 3 to achieve choice performance comparable to young rats. **C:** The mean indifference point (the delay at which rats showed equivalent preference for the small and large rewards) was significantly greater in aged rats compared to young. In all panels, error bars represent the standard error of the mean (SEM). *p<0.05, main effect of age; ^**^ p<0.01, age × delay block interaction.

### Effects of BLA Inactivation on choice behavior during deliberation

Inactivation of the BLA during the deliberation epoch (n=8 young and n=7 aged) significantly increased choice of the large reward to the same extent in young and aged rats, particularly at long delays (Figure 5A). A three-factor ANOVA (laser condition × age × delay block) indicated a main effect of laser condition (F_(1,13)_=103.507, p < 0.001, η^2^=0.888, 1-β=1.000) but no main effect of age (F_(1,13)_= 0.089, p=0.770, η^2^=0.007, 1-β=0.059) nor an age × laser condition interaction (F_(1,13)_=1.838, p = 0.198, η^2^=0.124, 1-β=0.242). A reliable main effect of delay block was observed (F_(2,26)_=112.005, p < 0.001, η^2^=0.896, 1-β=1.000), as was as an interaction between laser condition and delay block (F_(2,26)_=38.369, p < 0.001, η^2^=0.747, 1-β=1.000). Follow-up analyses, conducted to further explore the laser condition × delay block interaction, compared the effects of inactivation at each block. These analyses indicated that BLA inactivation significantly increased choice of the large reward in Blocks 2 (t_(14)_=−6.494, p<0.001, Cohen’s *d*=1.724, 1-β=0.995) and 3 (t_(14)_=−9.434, p<0.001, Cohen’s *d*=2.228, 1-β=1.000), but not in Block 1 in which rats of both ages strongly preferred the large reward, even under control conditions (t_(14)_=−0.323, p=0.751, Cohen’s *d*=0.124, 1-β=0.051).

**Figure 5.**
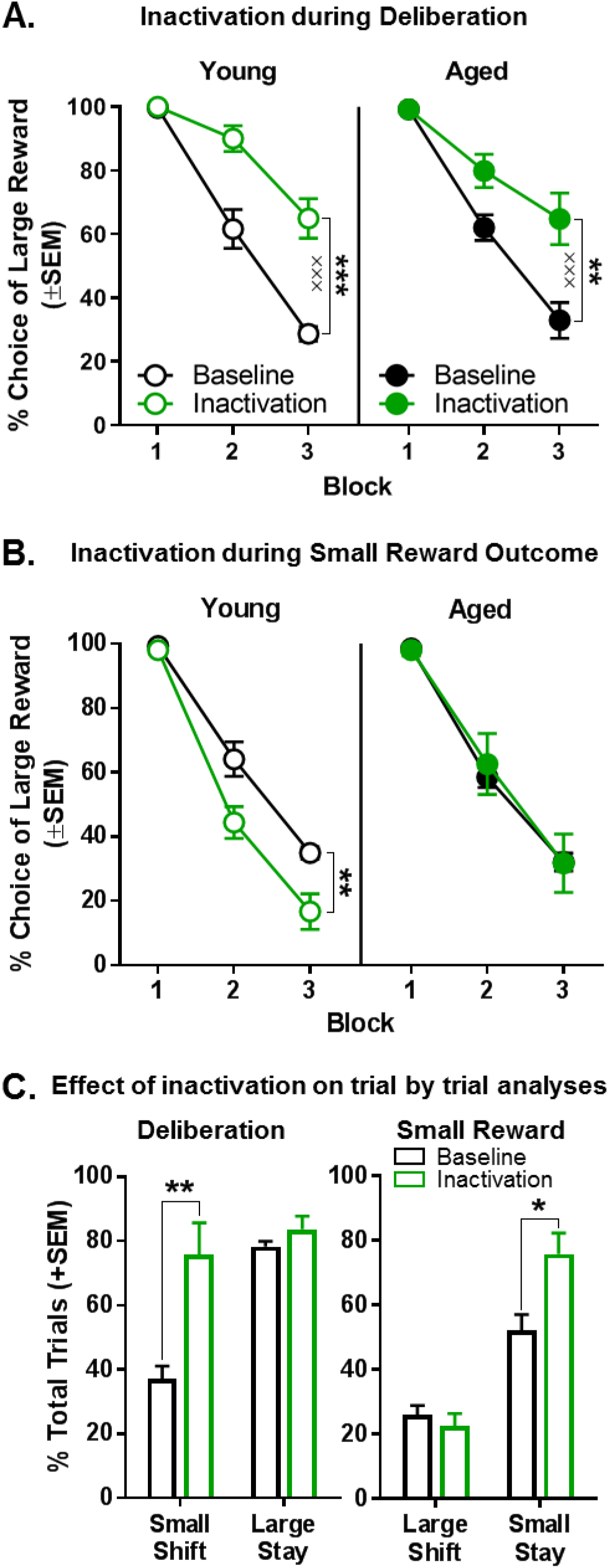
Effect of BLA inactivation during the deliberation and small reward epochs. **A:** Inactivation of the BLA during the deliberation epoch (prior to a choice) resulted in a significant increase in preference for the large, delayed reward in both young (n=8) and aged (n=7) rats. **B:** Inactivation of the BLA during the small reward epoch resulted in a significant decrease in preference for the large, delayed reward in young (n=6), but not aged (n=6), rats (**p<0.01, main effect of inactivation). **C:** Effects of BLA inactivation on trial-by-trial choice strategies in young rats. This analysis revealed that the increased choice of the large, delayed reward caused by BLA inactivation during deliberation in young rats (panel A) was due to an increase in the percentage of trials on which rats shifted to the large, delayed reward following a choice of the small, immediate reward. A similar analysis revealed that the decreased choice of the large, delayed reward caused by BLA inactivation during the small, immediate reward in young rats (panel B) was due to an increase in the percentage of trials on which rats “stayed” on the small, immediate reward following a choice of this reward on the previous trial. In all panels, error bars represent standard error of the mean (SEM). *p<.05, **p<0.01, ***p<0.001, main effect of inactivation; ^***^ p<0.001, inactivation × delay block interaction.

### Effects of BLA inactivation on choice behavior during the small reward

In direct contrast to the effects of BLA inactivation during deliberation, BLA inactivation during the small reward epoch (n=6 young and n=6 aged) significantly decreased choice of the large reward in young rats (Figure 5B). A three-factor ANOVA (laser condition × age × delay block) indicated main effects of laser condition (F_(1,10)_=5.131, p=0.047, η^2^=0.339, 1-β=0.534) and delay block (F_(2,20)_=248.854, p<0.001, η^2^=0.961, 1-β=1.000), but no interaction between laser condition and delay block (F_(2,20)_= 1.317; p = 0.290, η^2^=0.116, 1-β=0.251). Notably, although there was no main effect of age (F_(1,10)_=0.941; p=0.355, η^2^=0.086, 1-β=0.142), the effects of BLA inactivation during small reward delivery did reliably interact with age (age × inactivation condition: F_(1,10)_=7.127, p=0.024, η^2^=0.416, 1-β=0.673). To better define the nature of this interaction, two factor ANOVAs (laser condition × delay block) were performed separately on choice behavior in young and aged rats. BLA inactivation significantly decreased choice of the large reward in young rats (main effect of laser condition: F_(1,5)_=18.226; p=0.008, η^2^=0.785, 1-β=0.922, main effect of delay block: F_(2,10)_=173.588, p<0.001, η^2^=0.972, 1-β=1.000; laser condition × delay block: F_(2,10)_=3.829, p=0.058, η^2^=0.434, 1-β=0.556) but not in aged rats (main effect of laser condition: F_(1,5)_=0.061; p=0.814, η^2^=0.012, 1-β=0.055; main effect of delay block: F_(2,10)_=93.015, p<0.001, η^2^=0.949, 1-β=1.000; laser condition × delay block: F_(2,10)_=0.185, p=0.834, η^2^=0.036, 1-β=0.072).

### Altered choice strategy resulting from BLA inactivation during the deliberation and small reward epochs

The data above show that BLA inactivation in young rats during the deliberation and small reward epochs altered choice behavior in opposite directions (i.e., BLA inactivation during deliberation *increased* whereas BLA inactivation during small reward outcome *decreased* choice of the large, delayed reward). A trial-by-trial analysis was conducted on these data to determine the effects of BLA inactivation on two distinct behavioral strategies that could mediate these shifts in choice preference. Specifically, during the deliberation epoch, this analysis determined the degree to which BLA inactivation influenced rats to “shift” to the large reward option following a choice of the small reward on the previous trial, versus “stay” with the large reward option following a choice of large reward on the previous trial. In the small reward outcome epoch, the analysis assessed the degree to which BLA inactivation influenced rats to “shift” to the small reward option following a choice of the large reward on a previous trial, versus “stay” with the small reward option following a choice of the small reward on the previous trial.

As shown in Figure 5C, the percentage of trials during deliberation epoch inactivation on which a large reward choice was followed by a second large reward choice (large-stay) did not differ as a function of laser condition (t_(7)_=−1.299, p=0.235, Cohen’s *d*=0.618, 1-β=0.208). In contrast, a similar analysis conducted on the percentage of trials on which a choice of the small reward was followed by choice of the large reward (small-shift) revealed a main effect of laser condition (t_(7)_ =−4.095, p=0.005, Cohen’s *d*=1.802, 1-β=0.917). This finding indicates that the effects on choice behavior of BLA inactivation during deliberation result from rats shifting choices toward the large reward following a choice of the small reward. Applying a parallel analysis to sessions in which inactivation took place during the small reward epoch yielded a different pattern of results. BLA inactivation during the small reward epoch significantly increased the percentage of trials on which a small reward choice was followed by a second small reward choice (small-stay; t_(5)_=−3.593, p=0.016, Cohen’s *d*=1.694, 1-β=0.754). In contrast, BLA inactivation did not affect the percentage of trials on which a choice of the large reward was followed by a choice of the small reward (large-shift; t_(5)_=0.770, p=0.476, Cohen’s *d*=0.399, 1-β=0.091).

### Effects of BLA inactivation on other task performance measures during inactivation during the deliberation and small reward epochs

Other task-specific measures were compared between BLA inactivation and baseline conditions in both deliberation and small reward epochs. The number of trials completed did not differ as a function of laser condition or age in either epoch (see Table 1). Similarly, no differences in response latency were observed as a function BLA inactivation, age, or lever type in these epochs (See Table 2).

**Table 1.** Effects of BLA inactivation on number of trials completed. There were no effects of BLA inactivation during either the deliberation epoch (Fs_(1,13)_=0.162-3.264; ps=0.094-0.694) or the small reward epoch (Fs_(1,10)_=0.180-3.431; ps=0.094-0.681).

| Decision Making Period | Age | Inactivation Condition | Mean | Std. Error | N |
| --- | --- | --- | --- | --- | --- |
| Deliberation | Young | Baseline | 52.688 | 0.81 | 8 |
|  |  | Inactivation | 51.625 | 0.94 | 8 |
|  | Aged | Baseline | 51.714 | 0.87 | 7 |
|  |  | Inactivation | 53.429 | 1.003 | 7 |
| Small Reward Outcome | Young | Baseline | 52.667 | 0.394 | 6 |
|  |  | Inactivation | 53.667 | 0.635 | 6 |
|  | Aged | Baseline | 52.639 | 0.394 | 6 |
|  |  | Inactivation | 53.167 | 0.635 | 6 |

**Table 2.**
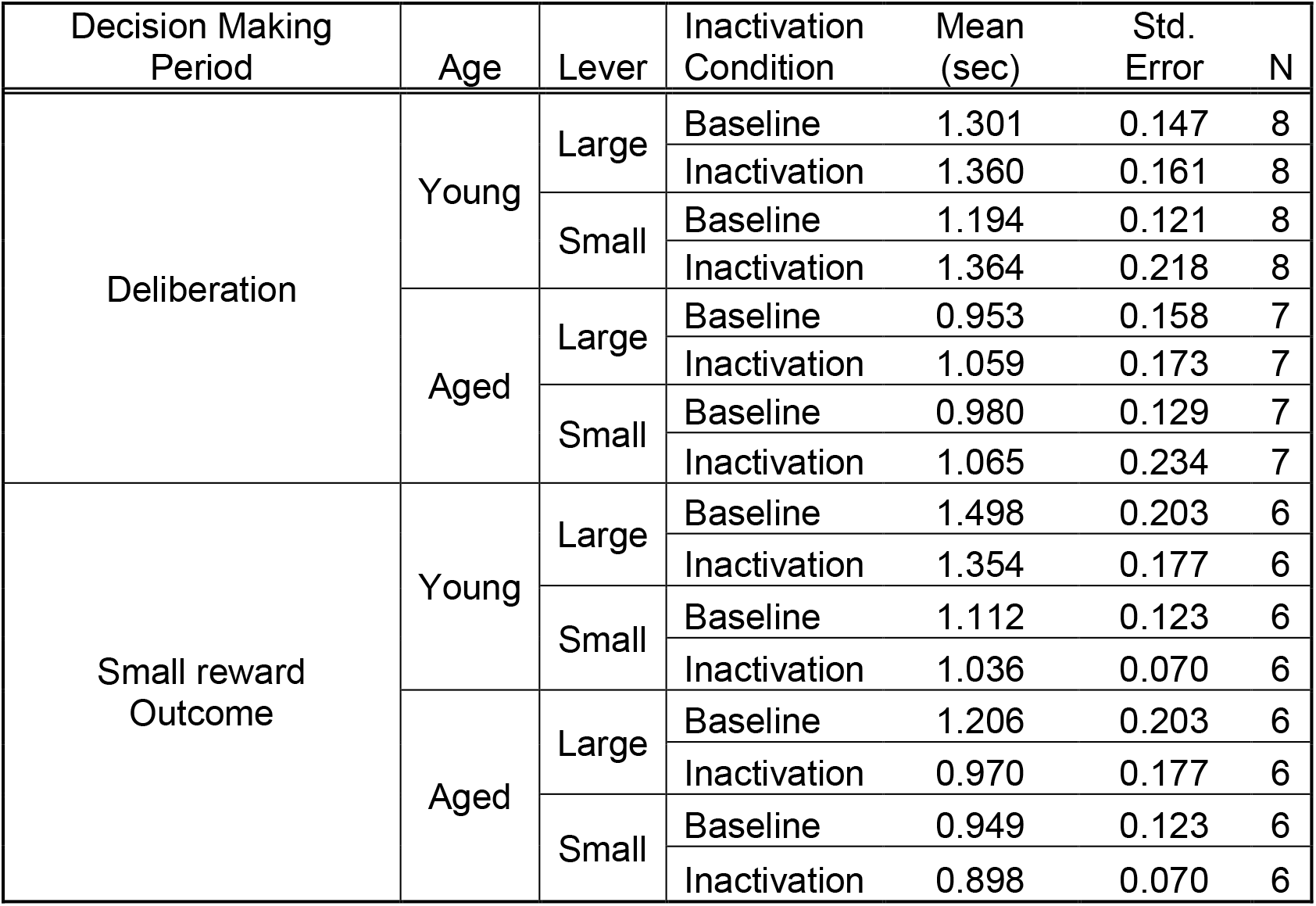
Effects of BLA inactivation on lever response latencies. There were no effects of BLA inactivation during either the deliberation epoch (Large reward lever: Fs_(1, 13)_=0.588-2.898, ps=0.112-0.457; Small reward lever: Fs_(1, 8)_=0.505-2.050, ps=0.190-0.497) or the small reward epoch (Large reward lever: Fs_(1, 10)_=0.257-4.149, ps=0.069-0.623; Small reward lever: Fs_(1, 10)_=0.004-3.225, ps=0.103-0.954).

| Decision Making Period | Age | Lever | Inactivation Condition | Mean (sec) | Std. Error | N |
| --- | --- | --- | --- | --- | --- | --- |
| Deliberation | Young | Large | Baseline | 1.301 | 0.147 | 8 |
|  |  |  | Inactivation | 1.360 | 0.161 | 8 |
|  |  | Small | Baseline | 1.194 | 0.121 | 8 |
|  |  |  | Inactivation | 1.364 | 0.218 | 8 |
|  | Aged | Large | Baseline | 0.953 | 0.158 | 7 |
|  |  |  | Inactivation | 1.059 | 0.173 | 7 |
|  |  | Small | Baseline | 0.980 | 0.129 | 7 |
|  |  |  | Inactivation | 1.065 | 0.234 | 7 |
| Small reward Outcome | Young | Large | Baseline | 1.498 | 0.203 | 6 |
|  |  |  | Inactivation | 1.354 | 0.177 | 6 |
|  |  | Small | Baseline | 1.112 | 0.123 | 6 |
|  |  |  | Inactivation | 1.036 | 0.070 | 6 |
|  | Aged | Large | Baseline | 1.206 | 0.203 | 6 |
|  |  |  | Inactivation | 0.970 | 0.177 | 6 |
|  |  | Small | Baseline | 0.949 | 0.123 | 6 |
|  |  |  | Inactivation | 0.898 | 0.070 | 6 |

### Effects of BLA inactivation during epochs associated with the large reward outcome

Choosing the large reward lever resulted in a variable delay period that was followed by large (3 food pellets) reward delivery. The effects of BLA inactivation during the delay and large reward delivery epochs were initially tested in separate sessions (n=6 young and n=6 aged). Subsequently, the effects of BLA inactivation across both the delay and large reward epochs were tested in a subset of these rats (n=3 young and n=3 aged).

#### Effects of BLA inactivation during the delay epoch

The effects of BLA inactivation during the delay epoch were tested in delay blocks 2 and 3 using a three-factor ANOVA (laser condition × age × block). As expected, there was a main effect of delay block (F_(2,20)_=146.811, p<0.001, η^2^=0.936, 1-β=1.000) such that both young and aged rats decreased their choice of the large reward as the delay prior to the large reward increased (Figure 6A). Compared to baseline, however, no reliable differences in choice behavior resulted from BLA inactivation during the delay epoch (F_(1,10)_=0.005, p=0.947, η^2^=0.000, 1-β=0.050), nor was there an interaction between inactivation condition and delay block (F_(2,20)_=0.002, p=0.998, η^2^=0.000, 1-β=0.050). Similarly, there were neither main effects nor interactions associated with age (main effect of age: F_(1,10)_<0.001, p=0.996, η^2^=0.000, 1-β=0.050; age × delay block: F_(2,20)_=0.077, p=0.926, η^2^=0.008, 1-β=0.060; age × laser condition: F_(1,10)_=0.081, p=0.782, η^2^=0.008, 1-β=0.058; age × laser condition × delay block: F_(2,20)_ = 0.096, p=0.908, η^2^=0.010, 1-β=0.063).

**Figure 6.**
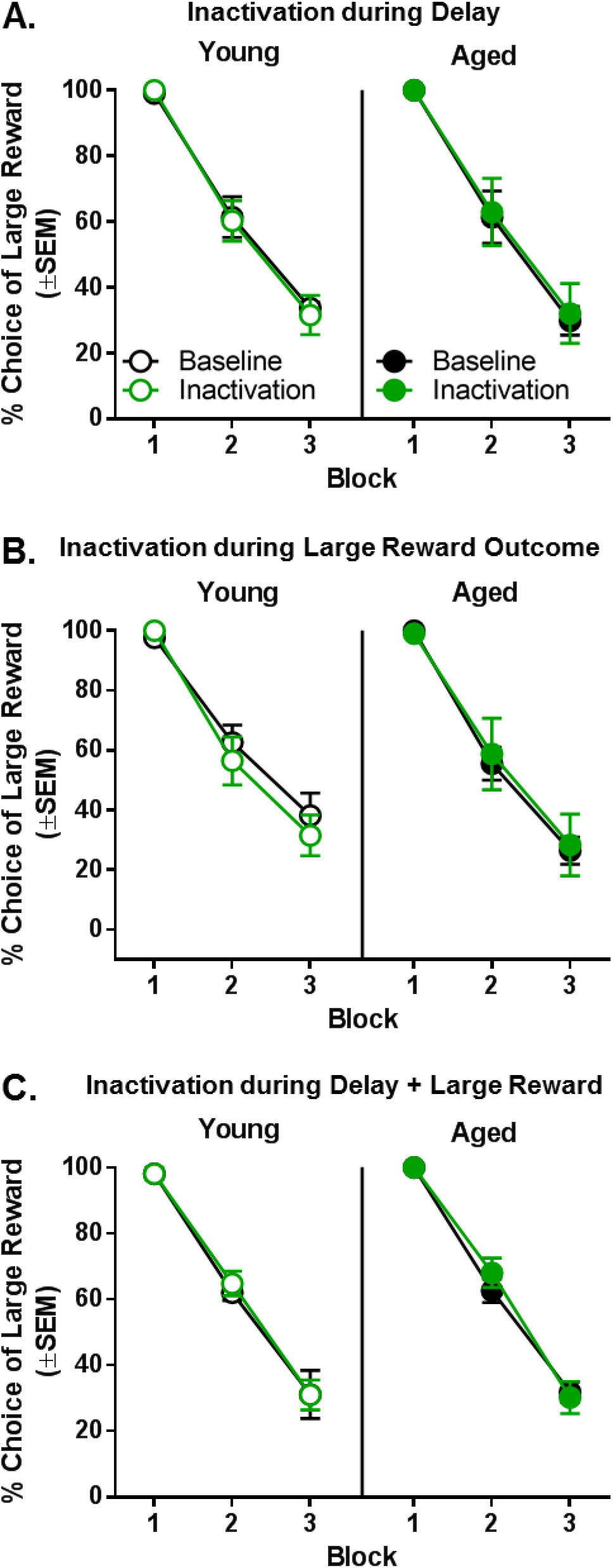
Effect of BLA inactivation during outcomes associated with choice of the large reward. **A:** Inactivation of the BLA during the delay epoch resulted in no change in choice performance in either young (n=6) or aged (n=6) rats. **B:** Inactivation of the BLA during the large reward epoch resulted in no change in choice performance in either young (n=6) or aged (n=6) rats. **C:** Inactivation of the BLA during both the delay and large reward epochs resulted in no change in choice performance in either young (n=3) or aged (n=3) rats. Error bars represent standard error of the mean (SEM).

#### Effects of BLA inactivation during the large reward epoch

Unlike the effects of BLA inactivation during the small reward epoch, inactivation of BLA during the large reward epoch did not alter choice performance in either young or aged rats compared to baseline (Figure 6B). As expected, there was a main effect of delay block (F_(2,20)_= 120.846, p<0.001, η^2^=0.924, 1-β=1.000) such that both young and aged rats decreased their choice of the large reward as the delay to large reward delivery increased. Compared to baseline, however, no reliable differences in choice behavior resulted from BLA inactivation during the large reward epoch (main effect of laser condition: F_(1,10)_=0.125, p=0.731, η^2^=0.012, 1-β=0.062), nor was there an interaction between laser condition and delay block (F_(2,20)_=0.133, p=0.876, η^2^=0.013, 1-β=0.068). Similarly, there were no main effects (F_(1,10)_=0.249, p=0.629, η^2^=0.024, 1-β=0.074) nor interactions associated with age (age × delay block: F_(2,20)_=0.437, p=0.652, η^2^=0.042, 1-β=0.111; age × laser condition: F_(1,10)_=0.697, p=0.423, η^2^=0.065, 1-β=0.118; age × laser condition × delay block: F_(2,20)_=0.664, p=0.526, η^2^=0.062, 1-β=0.146).

#### Effects of BLA inactivation during both delay and large reward epochs

One possible explanation for the null effects of BLA inactivation during either the delay or large reward epochs is that, given the role of the BLA in integration of rewards and costs, inactivation may only be effective when conducted during both of these epochs. To evaluate this possibility, rats were tested while the BLA was inactivated during *both* the delay and large reward epochs. Continuous inactivation across both epochs yielded no effects on choice performance. As shown in Figure 6C, a three-factor ANOVA (laser condition × age × delay block) revealed the expected main effect of delay block (F_(2,8)_=193.743, p<0.001, η^2^=0.980, 1-β=1.000) but no main effects or interactions with laser condition or age (main effect of laser condition: F_(1,4)_=0.757, p=0.433, η^2^=0.159, 1-β=0.105; main effect of age: F_(1,4)_=0.306, p=0.610, η^2^=0.071, 1-β=072; laser condition × delay block: F_(2,8)_=0.979, p=0.417, η^2^=0.197, 1-β=0.165; laser condition × age: F_(2,8)_=0.053, p=0.949, η^2^=0.006, 1-β=0.052; laser condition × age × delay block: F_(2,8)_=0.159, p=0.856, η^2^=0.038, 1-β=0.067).

### Effects of BLA inactivation during the intertrial interval

To confirm that BLA inactivation is not sufficient to produce effects on choice behavior in a non-temporally-specific manner, rats (n=6 young, n=6 aged) were tested while the BLA was inactivated during the intertrial interval (ITI). Although the expected main effect of delay block was observed (F_(2,20)_=116.459, p<0.001, η^2^=0.921, 1-β=1.000), BLA inactivation during the ITI did not alter choice performance compared to baseline in young or aged rats (main effect of laser condition: F_(1,10)_=0.082, p=0.780, η^2^=0.008, 1-β=0.058; main effect of age: F_(1,10)_=0.042, p=0.842, η^2^=0.004, 1-β=0.054; laser condition × age: F_(1,10)_=0.298, p=0.597, η^2^=0.029, 1-β=0.079; laser condition × delay block: F_(2,20)_=0.344, p=0.713, η^2^=0.033, 1-β=0.097; age × delay block: F_(2,20)_=0.216, p=0.808, η^2^=0.021, 1-β=0.079; age × laser condition × delay block: F_(2,20)_=0.198; p=0.822, η^2^=0.019, 1-β=0.077; Figure 7).

**Figure 7.**
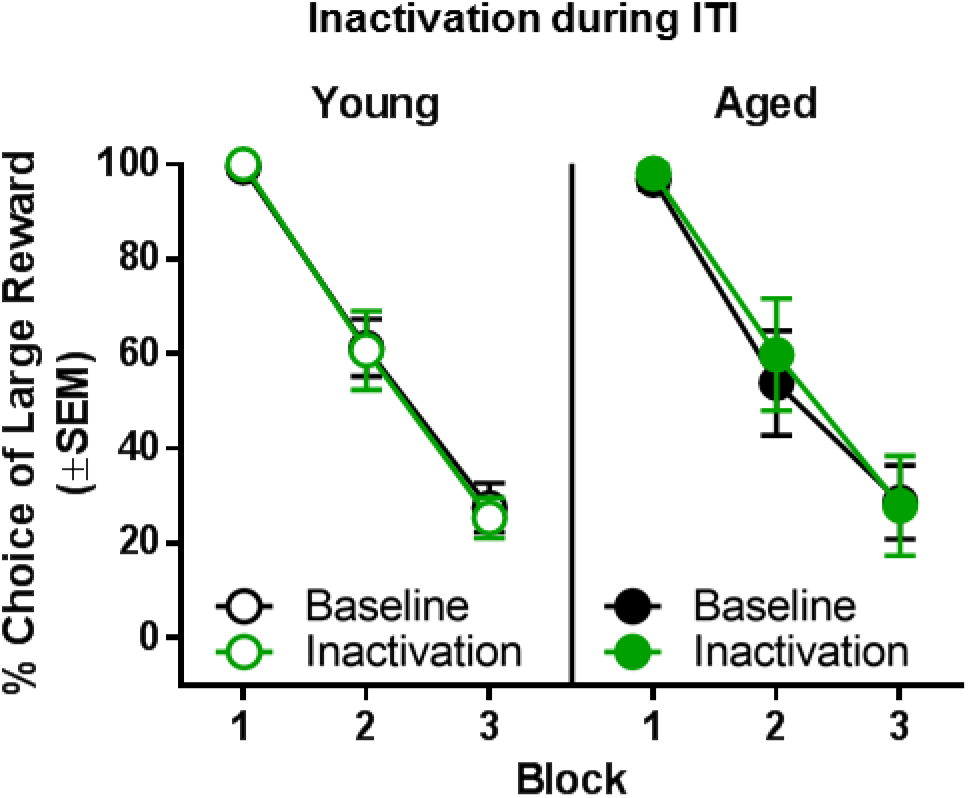
Effect of BLA inactivation during the intertrial interval. Inactivation of the BLA during the intertrial interval resulted in no change in choice performance in either young (n=6) or aged (n=6) rats. Error bars represent standard error of the mean (SEM).

### Effects of light delivery into BLA in rats with control virus (AAV5-CamkIIα-mCherry)

To control for non-specific effects of light delivery (e.g., changes in tissue temperature), the effects of light delivery in rats transduced with a control virus that did not contain the eNpHR3.0 gene were tested during behavioral epochs in which BLA inactivation influenced choice behavior [i.e, deliberation (n=4 young and n=4 aged rats) and small reward (n=4 young rats)].

#### Effects of light delivery during the deliberation epoch

Light delivery during the deliberation epoch in rats transduced with a control virus had no effects on choice performance (Figure 8A). A three factor ANOVA (laser condition × age × delay block) indicated the expected main effect of delay block (F_(2,12)_=100.272; p<0.001, η^2^=0.944, 1-β=1.000) but no main effects or interactions involving laser condition or age (main effect of laser condition: F_(1,6)_=0.128; p=0.733, η^2^=0.021, 1-β=0.061; main effect of age: F_(1,6)_=0.055; p=0.823, η^2^=0.009, 1-β=0.055; laser condition × age: F_(1,6)_=0.028; p=0.874, η^2^=0.005, 1-β=0.052; laser condition × delay block: F_(2,12)_=0.121; p=0.887, η^2^=0.020, 1-β=0.065; age × delay block: F_(2,12)_=0.105; p=0.902;, η^2^=0.017, 1-β=0.063 laser condition × age × delay block: F_(2,12)_=0.434; p=0.658, η^2^=0.067, 1-β=0.105).

**Figure 8.**
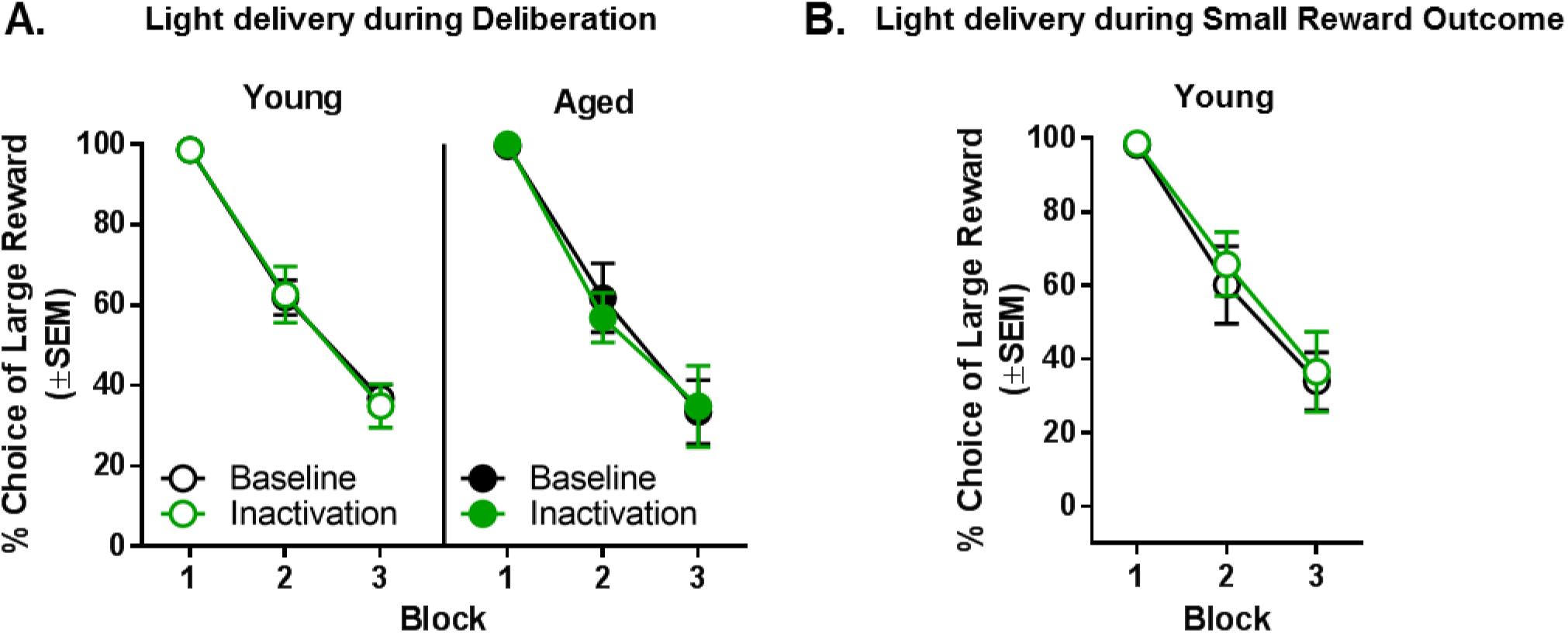
Effect of light delivery into BLA during the deliberation and small reward e in rats transduced with a control vector. **A:** Light delivery into the BLA during the deliberation epoch resulted in no change in choice performance in either young (n=4) or aged (n=4) control vector rats. **C:** Light delivery into the BLA during the small reward epoch resulted in no change in choice performance in young (n=4) control vector rats. Error bars represent standard error of the mean (SEM).

#### Effects of light delivery during the small reward epoch

Light delivery during the small reward epoch in young rats transduced with control virus also failed to influence choice performance (Figure 8B). A two factor ANOVA (laser condition × delay block) indicated the expected effect of delay block (F_(2,6)_=46.712; p<0.001, η^2^=0.940, 1-β=1.000) but no main effect of laser condition (F_(1,3)_=0.359; p=0.592, η^2^=0.107, 1-β=0.072) or laser condition × delay block interaction (F_(2,6)_=0.173; p=0.845, η^2^=0.055, 1-β=0.067).

## Discussion

### Age Differences in Intertemporal Choice

Across species, aging is accompanied by increased preference for large, delayed over small, immediate rewards (Green et al., 1994, 1999; Simon et al., 2010; Jimura et al., 2011; Löckenhoff et al., 2011; Samanez-Larkin et al., 2011; Eppinger et al., 2012; Hernandez et al., 2017). Previous work from our labs showed that relative to young rats, aged rats display greater preference for large, delayed over small, immediate rewards in a “fixed delays, block design” intertemporal choice task. This difference is not readily attributable to age-related deficits in cognitive flexibility, working memory, or food motivation, nor is it attributable to impairments in reward or temporal discrimination (Simon et al., 2010; Hernandez et al., 2017). The present study replicated these prior findings using a task variant in which the fixed delays/block design used in our previous work was maintained, but the delays to large reward delivery were adjusted on an individual basis to obtain equivalent levels of choice preference in young and aged rats. Under these conditions, aged rats required longer delays to achieve levels of choice preference comparable to young, suggesting that delays are less effective at discounting reward value in aged compared to young rats. These data are consistent with findings in human subjects (Green et al., 1994, 1999; Jimura et al., 2011; Eppinger et al., 2012) and indicate that an enhanced ability to delay gratification is a consistent feature of aging across species.

Data from the current study leveraged optogenetic approaches in young and aged rats to elucidate the contributions of BLA to intertemporal choice in young adult rats and to age-associated changes in this aspect of decision making. Temporally-discrete inactivation revealed distinct roles for BLA in intertemporal choice during the periods immediately before and after a choice was made. Specifically, BLA inactivation during the period prior to a choice (the deliberation epoch) increased choice of large, delayed over small, immediate rewards in both young and aged rats. In contrast, BLA inactivation during receipt of the small, immediate reward (after the choice) decreased choice of the large, delayed reward in young rats, whereas the same inactivation in aged rats had no effect on choice behavior. It should be noted that these results were not likely due to non-specific effects on behavior. Light delivery in rats expressing the control vector did not affect choice behavior, demonstrating that effects of BLA inactivation are due specifically to optogenetic rather than non-specific mechanisms. The fact that inactivation during the delay, large reward delivery, and ITI epochs had no effect on choice performance lends further specificity to the effects of BLA inactivation. Moreover, there were no effects of BLA inactivation during either the deliberation or small reward epochs on the number of trials completed or response latencies. Finally, effects on choice performance were not driven by alterations in reward magnitude discrimination, as BLA inactivation during either the deliberation or small reward epochs did not alter choice performance under no delay conditions (Block 1). The fact that effects were observed only under conditions in which a cost accompanied the large reward (Blocks 2 and 3) suggests a role for the BLA in assigning value to rewards based on cost parameters. Considered together, these data indicate that BLA activity is engaged in different roles at distinct points in the decision process, and that these roles change across the lifespan.

### BLA and Reward Outcome Evaluation

BLA inactivation during the small reward epoch decreased choice of large, delayed rewards in young rats, concurrent with a selective increase in repetitive choices of small, immediate rewards on the trial-by-trial analysis. This pattern of results, which is similar to that induced by less temporally-specific approaches to inhibiting BLA activity, suggests that BLA inactivation causes a failure to acquire or integrate information about the negative properties of the small reward (i.e., that it is smaller than the large reward), rendering rats more likely to choose this option on subsequent trials. This idea is consistent with a large literature supporting a critical role for BLA in behavior directed by reward value (i.e., behavior that is sensitive to shifts in reward value; Hatfield et al., 1996; Málková et al., 1997; Baxter et al., 2000; Shiflett and Balleine, 2010; Izquierdo et al., 2013; Parkes and Balleine, 2013). Moreover, these results are consistent with the effects of BLA inactivation during receipt of a large, punished reward in a risky decision making task. Inactivation of the BLA caused an increase in choice of this option, suggesting that under normal circumstances, this structure processes information about the negative qualities of outcomes to bias future behavior toward more favorable options (Orsini et al., 2017).

The finding that BLA inactivation during the small reward epoch (which decreased choice of large, delayed rewards in young rats) had no effect in aged rats indicates that aging is associated with a reduced role for BLA in reward outcome evaluation. Importantly, this lack of effect in aged rats was likely not due to age-related impairments in viral transduction or optogenetic efficacy, as BLA inactivation during the deliberation epoch in these same aged rats produced effects on behavior that were as robust as in young rats. In addition, because delays were adjusted to equate baseline performance in young and aged rats, the lack of effect in aged rats cannot be attributed to insufficient parametric space. Instead, the body of data in the current study support the idea of multiple BLA circuits that play unique roles in intertemporal decision making, and indicate that these circuits are differentially recruited in aged subjects. BLA neurons respond differentially to reward anticipation and specific outcomes (Schoenbaum et al., 1998; Belova et al., 2007, 2008; Sangha et al., 2013, 2013; Zhang et al., 2013; Beyeler et al., 2016), and distinct BLA efferents are thought to mediate encoding of appetitive vs. aversive stimuli (Beyeler et al., 2016; Burgos-Robles et al., 2017). Similar principles may operate in the context of intertemporal choice. Specifically, with respect to reward outcome evaluation, BLA projections to the nucleus accumbens (NAc) and other striatal regions may be particularly relevant. Pharmacological disconnection of the BLA and NAc impairs discrimination between a devalued vs. a non-devalued food reward (Shiflett and Balleine, 2010), and a recent study in rats performing a probabilistic decision making task (choices between a small, guaranteed reward vs. a large, probabilistic reward) showed that optogenetic inactivation of BLA terminals in NAc during reward omission caused an increase in preference for this “risky” option (Bercovici et al., 2018). These results support the idea that during intertemporal choice, activity in a BLA-NAc circuit integrates information about reward outcomes in order to shift future decisions toward choice of options associated with larger but delayed rewards by using feedback about the value (or lack thereof) of the smaller, more immediate reward. Notably, Eppinger et al. (2013) showed blunted activity in ventral striatum during reward prediction errors in older adults performing a learning task. Moreover, with respect to BLA circuits, a recent study by Samson et al. (2017) reported enhanced β-power in BLA during evaluation of reward in a probabilistic decision making task, which may reflect restructuring of reward networks during aging. Indeed, in both humans and rats, there is substantial evidence for recruitment of brain circuits that are distinct from those engaged by young subjects performing complex cognitive operations, even when equated for performance (Antonenko and Flöel, 2014; Lighthall et al., 2014; Tomás-Pereira et al., 2015; Wang et al., 2015). It has further been suggested that in comparison to young, aged subjects use different information to make decisions, relying more heavily on compensatory cognitive strategies and differentially weighting rewards and costs (Löckenhoff et al., 2011; Mather et al., 2012; Samanez-Larkin and Knutson, 2015; Pachur et al., 2017). These distinct cognitive strategies in older adults may minimize the role of reward outcome evaluation in guiding intertemporal choices.

### BLA and Deliberation

Unlike inactivation during outcome evaluation, optogenetic inhibition of BLA during deliberation in both young and aged rats closely mimicked the effects of age on intertemporal choice (i.e., increased choice of the large, delayed reward; Figure 4C and Simon et al., 2010; Hernandez et al., 2017). This role for the BLA in choice behavior is only observed using temporally-discrete optogenetic inhibition during deliberation, and not with experimental methods such as lesions or pharmacological inactivation that inhibit the BLA across all stages of the decision process and which produce opposite effects on choice behavior (i.e., decreased choice of large, delayed rewards; Winstanley et al., 2004; Churchwell et al., 2009). The fact that BLA circuits involved in reward outcome evaluation are not engaged in aged subjects thus may “unmask” the influence of a putative “BLA-deliberation” circuit on choice behavior. This pattern of results observed following BLA inactivation during deliberation in intertemporal choice is similar to those in a prior study from our labs which showed that BLA inactivation during the deliberation epoch in a risky decision making task decreased preference for risky rewards (Orsini et al., 2017). One interpretation of the finding that BLA activity during deliberation contributes to both impulsive and risky choices is that this activity is important for incentive motivation, driving individuals to seek more immediate rewards during intertemporal choices and larger, more salient rewards despite potential punishment during risky choices. This interpretation agrees with evidence from other behavioral contexts. For example, an intact BLA is necessary for the potentiating influence of reward-predictive cues on instrumental responding for reward (Everitt et al., 2003), as well as for maintenance of effortful choices of preferred options (Hart and Izquierdo, 2017). In addition, the trial-by-trial analysis of the current data indicates that BLA inactivation during deliberation increases the frequency with which rats shift from choosing the small, immediate to the large, delayed reward.

In addition to its projections to the NAc described above, the BLA projects to many other output structures (Sah et al., 2003), some of which may play a unique role in guiding intertemporal choices. In particular, BLA projections to PFC may confer incentive information about potential outcomes prior to the choice (Pérez-Jaranay and Vives, 1991; St Onge and Floresco, 2010; Sripada et al., 2011; Dilgen et al., 2013; Kim et al., 2017). The existence of such a putative BLA-PFC “deliberation circuit” is supported by recent data showing that neural activity in PFC is preceded by activity in BLA in response to conditioned reward cues (Burgos-Robles et al., 2017). While under normal conditions, activity in BLA circuits involved in outcome evaluation may be the primary driver of choice behavior, the failure to engage such circuits in aging may shift the influence of BLA to those circuits engaged prior to a choice, during deliberation of the choice options. As such, structural or functional changes in BLA that occur with aging are likely to exert their influence on intertemporal choice through this putative “deliberation circuit”. Indeed, studies in humans and rodents have shown that neural activity in anticipation of both rewarding and aversive stimuli can be blunted in older subjects compared to young (Schoenbaum et al., 2006; Eppinger et al., 2015; Samanez-Larkin and Knutson, 2015). Further, electrophysiological data indicate that BLA activity, as assessed by baseline firing rate of BLA neurons *in vivo*, is reduced in aged rats (Roesch et al., 2012). These data, together with those in the current study, suggest that BLA circuits that normally encode the incentive value of rewards may be hypoactive in aging. In combination with a failure to engage BLA circuits during outcome evaluation, these effects of aging on BLA neurons may contribute to the attenuated impulsive choice observed in aging. Experiments focusing on specific molecular mechanisms underlying age differences in neural activity within discrete BLA circuits will be useful for elucidating the neural substrates that account for the increased ability of aged subjects to delay gratification. An increased appreciation of such mechanisms within the context of the nuanced roles of BLA across multiple stages of the decision process could reveal therapeutic targets for optimizing decision making in both older and younger adults.

## Acknowledgements

We thank Vicky S. Kelly, Shannon C. Wall, and Bonnie I. McLaurin for technical assistance. Supported by R01AG029421 and the McKnight Brain Research Foundation (JLB), RF1AG060778 (JLB, BS, CJF), a McKnight Predoctoral Fellowship and the Pat Tillman Foundation (CMH), and a Thomas H. Maren Fellowship and K99DA041493 (CAO).

## Abbreviations

(BLA): basolateral amygdala
(FBN): Fischer 344 x Brown Norway hybrid
(ITI): intertrial interval

## References

Antonenko D, Flöel A (2014) Healthy Aging by Staying Selectively Connected: A Mini-Review. Gerontology 60:3–9.

Bailey MR, Simpson EH, Balsam PD (2016) Neural substrates underlying effort, time, and risk-based decision making in motivated behavior. Neurobiol Learn Mem 133:233–256.

Baxter MG, Parker A, Lindner CCC, Izquierdo AD, Murray EA (2000) Control of Response Selection by Reinforcer Value Requires Interaction of Amygdala and Orbital Prefrontal Cortex. J Neurosci 20:4311–4319.

Beas BS, Setlow B, Gregory R, Samanez-Larkin, Bizon JL (2015) Modeling Cost-Benefit Decision Making in Aged Rodents. In: Aging and Decision Making: Empirical and Applied Perspectives, pp 16–40. Elsevier.

Belova MA, Paton JJ, Morrison SE, Salzman CD (2007) Expectation Modulates Neural Responses to Pleasant and Aversive Stimuli in Primate Amygdala. Neuron 55:970–984.

Belova MA, Paton JJ, Salzman CD (2008) Moment-to-moment tracking of state value in the amygdala. J Neurosci Off J Soc Neurosci 28:10023–10030.

Bercovici DA, Princz-Lebel O, Tse MT, Moorman DE, Floresco SB (2018) Optogenetic Dissection of Temporal Dynamics of Amygdala-Striatal Interplay during Risk/Reward Decision Making. eNeuro 5:ENEURO.0422-18.2018.

Beyeler A, Namburi P, Glober GF, Simonnet C, Calhoon GG, Conyers GF, Luck R, Wildes CP, Tye KM (2016) Divergent Routing of Positive and Negative Information from the Amygdala during Memory Retrieval. Neuron 90:348–361.

Bickel WK, Koffarnus MN, Moody L, Wilson AG (2014) The behavioral- and neuro-economic process of temporal discounting: A candidate behavioral marker of addiction. Neuropharmacology 76:518–527.

Boyle PA, Yu L, Wilson RS, Gamble K, Buchman AS, Bennett DA (2012) Poor Decision Making Is a Consequence of Cognitive Decline among Older Persons without Alzheimer’s Disease or Mild Cognitive Impairment. PLOS ONE 7:e43647.

Burgos-Robles A, Kimchi EY, Izadmehr EM, Porzenheim MJ, Ramos-Guasp WA, Nieh EH, Felix-Ortiz AC, Namburi P, Leppla CA, Presbrey KN, Anandalingam KK, Pagan-Rivera PA, Anahtar M, Beyeler A, Tye KM (2017) Amygdala inputs to prefrontal cortex guide behavior amid conflicting cues of reward and punishment. Nat Neurosci 20:824–835.

Burke SN, Thome A, Plange K, Engle JR, Trouard TP, Gothard KM, Barnes CA (2014) Orbitofrontal Cortex Volume in Area 11/13 Predicts Reward Devaluation, But Not Reversal Learning Performance, in Young and Aged Monkeys. J Neurosci 34:9905–9916.

Churchwell JC, Morris AM, Heurtelou NM, Kesner RP (2009) Interactions between the prefrontal cortex and amygdala during delay discounting and reversal. Behav Neurosci 123:1185–1196.

Crowley TJ, Dalwani MS, Sakai JT, Raymond KM, McWilliams SK, Banich MT, Mikulich-Gilbertson SK (2017) Children’s brain activation during risky decision-making: A contributor to substance problems? Drug Alcohol Depend 178:57–65.

Decker JH, Figner B, Steinglass JE (2015) On Weight and Waiting: Delay Discounting in Anorexia Nervosa Pretreatment and Posttreatment. Biol Psychiatry 78:606–614.

Denburg NL, Cole CA, Hernandez M, Yamada TH, Tranel D, Bechara A, Wallace RB (2007) The Orbitofrontal Cortex, Real-World Decision Making, and Normal Aging. Ann N Y Acad Sci 1121:480–498.

Dilgen J, Tejeda HA, O’Donnell P (2013) Amygdala inputs drive feedforward inhibition in the medial prefrontal cortex. J Neurophysiol 110:221–229.

Eppinger B, Nystrom LE, Cohen JD (2012) Reduced sensitivity to immediate reward during decision-making in older than younger adults. PloS One 7:e36953.

Eppinger B, Schuck NW, Nystrom LE, Cohen JD (2013) Reduced striatal responses to reward prediction errors in older compared with younger adults. J Neurosci Off J Soc Neurosci 33:9905–9912.

Eppinger B, Heekeren HR, Li SC (2015) Age-related prefrontal impairments implicate deficient prediction of future reward in older adults. Neurobiol Aging 36:2380–2390.

Evenden JL, Ryan CN (1996) The pharmacology of impulsive behaviour in rats: the effects of drugs on response choice with varying delays of reinforcement. Psychopharmacology (Berl) 128:161–170.

Everitt BJ, Cardinal RN, Parkinson JA, Robbins TW (2003) Appetitive behavior: impact of amygdala-dependent mechanisms of emotional learning. Ann N Y Acad Sci 985:233–250.

Fobbs WC, Mizumori SJY (2017) A framework for understanding and advancing intertemporal choice research using rodent models. Neurobiol Learn Mem 139:89–97.

Ghods-Sharifi S, St Onge JR, Floresco SB (2009) Fundamental contribution by the basolateral amygdala to different forms of decision making. J Neurosci Off J Soc Neurosci 29:5251–5259.

Grabenhorst F, Hernádi I, Schultz W (2012) Prediction of economic choice by primate amygdala neurons. Proc Natl Acad Sci:201212706.

Green L, Fry AF, Myerson J (1994) Discounting of Delayed Rewards: A Life-Span Comparison. Psychol Sci 5:33–36.

Green L, Myerson J, Lichtman D, Rosen S, Fry A (1996) Temporal discounting in choice between delayed rewards: the role of age and income. Psychol Aging 11:79–84.

Green L, Myerson J, Ostaszewski P (1999) Discounting of delayed rewards across the life span: age differences in individual discounting functions. Behav Processes 46:89–96.

Hamilton KR et al. (2015) Choice Impulsivity: Definitions, Measurement Issues, and Clinical Implications. Personal Disord 6:182–198.

Hart EE, Izquierdo A (2017) Basolateral amygdala supports the maintenance of value and effortful choice of a preferred option. Eur J Neurosci 45:388–397.

Hatfield T, Han J-S, Conley M, Gallagher M, Holland P (1996) Neurotoxic Lesions of Basolateral, But Not Central, Amygdala Interfere with Pavlovian Second-Order Conditioning and Reinforcer Devaluation Effects. J Neurosci 16:5256–5265.

Hernandez CM, Vetere LM, Orsini CA, McQuail JA, Maurer AP, Burke SN, Setlow B, Bizon JL (2017) Decline of prefrontal cortical-mediated executive functions but attenuated delay discounting in aged Fischer 344 × brown Norway hybrid rats. Neurobiol Aging 60:141–152.

Hess TM, Strough J, Lockenhoff CE (2015) “Aging and decision making; empirical and applied perspectives.” Academic Press, San Diego, CA. 399 pages.

Izquierdo A, Darling C, Manos N, Pozos H, Kim C, Ostrander S, Cazares V, Stepp H, Rudebeck PH (2013) Basolateral Amygdala Lesions Facilitate Reward Choices after Negative Feedback in Rats. J Neurosci 33:4105–4109.

Jimura K, Myerson J, Hilgard J, Keighley J, Braver TS, Green L (2011) Domain independence and stability in young and older adults’ discounting of delayed rewards. Behav Processes 87:253–259.

Johansen JP, Wolff SBE, Lüthi A, LeDoux JE (2012) Controlling the Elements: An Optogenetic Approach to Understanding the Neural Circuits of Fear. Biol Psychiatry 71:1053–1060.

Kaye WH, Wierenga CE, Bailer UF, Simmons AN, Bischoff-Grethe A (2013) Nothing tastes as good as skinny feels: the neurobiology of anorexia nervosa. Trends Neurosci 36:110–120.

Kim CK, Ye L, Jennings JH, Pichamoorthy N, Tang DD, Yoo A-CW, Ramakrishnan C, Deisseroth K (2017) Molecular and Circuit-Dynamical Identification of Top-Down Neural Mechanisms for Restraint of Reward Seeking. Cell 170:1013–1027.e14.

Lighthall NR, Huettel SA, Cabeza R (2014) Functional Compensation in the Ventromedial Prefrontal Cortex Improves Memory-Dependent Decisions in Older Adults. J Neurosci 34:15648–15657.

Löckenhoff CE, O’Donoghue T, Dunning D (2011) Age differences in temporal discounting: the role of dispositional affect and anticipated emotions. Psychol Aging 26:274–284.

Lolova I, Davidoff M (1991) Changes in GABA-immunoreactivity and GABA-transaminase activity in rat amygdaloid complex in aging. J Hirnforsch 32:231–238.

Málková L, Gaffan D, Murray EA (1997) Excitotoxic Lesions of the Amygdala Fail to Produce Impairment in Visual Learning for Auditory Secondary Reinforcement But Interfere with Reinforcer Devaluation Effects in Rhesus Monkeys. J Neurosci 17:6011–6020.

Mata R, Josef AK, Samanez-Larkin GR, Hertwig R (2011) Age differences in risky choice: a meta-analysis. Ann N Y Acad Sci 1235:18–29.

Mather M, Mazar N, Gorlick MA, Lighthall NR, Burgeno J, Schoeke A, Ariely D (2012) Risk preferences and aging: the “certainty effect” in older adults’ decision making. Psychol Aging 27:801–816.

Orsini CA, Hernandez CM, Singhal S, Kelly KB, Frazier CJ, Bizon JL, Setlow B (2017) Optogenetic Inhibition Reveals Distinct Roles for Basolateral Amygdala Activity at Discrete Time Points during Risky Decision Making. J Neurosci 37:11537–11548.

Orsini CA, Moorman DE, Young JW, Setlow B, Floresco SB (2015a) Neural mechanisms regulating different forms of risk-related decision-making: Insights from animal models. Neurosci Biobehav Rev 58:147–167.

Orsini CA, Trotta RT, Bizon JL, Setlow B (2015b) Dissociable roles for the basolateral amygdala and orbitofrontal cortex in decision-making under risk of punishment. J Neurosci Off J Soc Neurosci 35:1368–1379.

Pachur T, Mata R, Hertwig R (2017) Who Dares, Who Errs? Disentangling Cognitive and Motivational Roots of Age Differences in Decisions Under Risk. Psychol Sci 28:504–518.

Parkes SL, Balleine BW (2013) Incentive Memory: Evidence the Basolateral Amygdala Encodes and the Insular Cortex Retrieves Outcome Values to Guide Choice between Goal-Directed Actions. J Neurosci 33:8753–8763.

Patros CHG, Alderson RM, Kasper LJ, Tarle SJ, Lea SE, Hudec KL (2016) Choice-impulsivity in children and adolescents with attention-deficit/hyperactivity disorder (ADHD): A meta-analytic review. Clin Psychol Rev 43:162–174.

Paxinos G, Watson C (2005) The rat brain in stereotaxic coordinates. New York: Elsevier Academic.

Pérez-Jaranay JM, Vives F (1991) Electrophysiological study of the response of medial prefrontal cortex neurons to stimulation of the basolateral nucleus of the amygdala in the rat. Brain Res 564:97–101.

Peters J, Büchel C (2011) The neural mechanisms of inter-temporal decision-making: understanding variability. Trends Cogn Sci 15:227–239.

Prager EM, Bergstrom HC, Wynn GH, Braga MFM (2016) The basolateral amygdala γ-aminobutyric acidergic system in health and disease. J Neurosci Res 94:548–567.

Rangel A, Camerer C, Montague PR (2008) A framework for studying the neurobiology of value-based decision making. Nat Rev Neurosci 9:545–556.

Roesch MR, Esber GR, Bryden DW, Cerri DH, Haney ZR, Schoenbaum G (2012) Normal aging alters learning and attention-related teaching signals in basolateral amygdala. J Neurosci Off J Soc Neurosci 32:13137–13144.

Rubinow MJ, Drogos LL, Juraska JM (2009) Age-related dendritic hypertrophy and sexual dimorphism in rat basolateral amygdala. Neurobiol Aging 30:137–146.

Rubinow MJ, Juraska JM (2009) Neuron and glia numbers in the basolateral nucleus of the amygdala from preweaning through old age in male and female rats: A stereological study. J Comp Neurol 512:717–725.

Sah P, Faber ESL, Armentia MLD, Power J (2003) The Amygdaloid Complex: Anatomy and Physiology. Physiol Rev 83:803–834.

Samanez-Larkin GR, Knutson B (2015) Decision making in the ageing brain: changes in affective and motivational circuits. Nat Rev Neurosci 16:278–289.

Samanez-Larkin GR, Mata R, Radu PT, Ballard IC, Carstensen LL, McClure SM (2011) Age Differences in Striatal Delay Sensitivity during Intertemporal Choice in Healthy Adults. Front Neurosci 5:126.

Samson RD, Lester AW, Duarte L, Venkatesh A, Barnes CA (2017) Emergence of β-Band Oscillations in the Aged Rat Amygdala during Discrimination Learning and Decision Making Tasks. eNeuro 4.

Sangha S, Chadick JZ, Janak PH (2013) Safety Encoding in the Basal Amygdala. J Neurosci 33:3744–3751.

Schoenbaum G, Chiba AA, Gallagher M (1998) Orbitofrontal cortex and basolateral amygdala encode expected outcomes during learning. Nat Neurosci 1:155–159.

Schoenbaum G, Chiba AA, Gallagher M (1999) Neural Encoding in Orbitofrontal Cortex and Basolateral Amygdala during Olfactory Discrimination Learning. J Neurosci 19:1876–1884.

Schoenbaum G, Setlow B, Saddoris MP, Gallagher M (2006) Encoding changes in orbitofrontal cortex in reversal-impaired rats. J Neurophysiol 95: 1509–1517.

Shiflett MW, Balleine BW (2010) At the limbic–motor interface: disconnection of basolateral amygdala from nucleus accumbens core and shell reveals dissociable components of incentive motivation. Eur J Neurosci 32:1735–1743.

Simon NW, LaSarge CL, Montgomery KS, Williams MT, Mendez IA, Setlow B, Bizon JL (2010) Good things come to those who wait: attenuated discounting of delayed rewards in aged Fischer 344 rats. Neurobiol Aging 31:853–862.

Sripada CS, Gonzalez R, Phan KL, Liberzon I (2011) The neural correlates of intertemporal decision-making: Contributions of subjective value, stimulus type, and trait impulsivity. Hum Brain Mapp 32:1637–1648.

St Onge JR, Floresco SB (2010) Prefrontal cortical contribution to risk-based decision making. Cereb Cortex N Y N 1991 20:1816–1828.

Steinglass JE, Figner B, Berkowitz S, Simpson HB, Weber EU, Walsh BT (2012) Increased Capacity to Delay Reward in Anorexia Nervosa. J Int Neuropsychol Soc 18:773–780.

Tomas-Pereira I, Gallagher M, Rapp PR (2015) Head west or left, east or right: interactions between memory systems in neurocognitive aging. Neurobiol Aging 36:3067–3078.

Wang W-C, Dew ITZ, Cabeza R (2015) Age-related differences in medial temporal lobe involvement during conceptual fluency. Brain Res 1612:48–58.

Wassum KM, Izquierdo A (2015) The basolateral amygdala in reward learning and addiction. Neurosci Biobehav Rev 57:271–283.

Winstanley CA, Theobald DEH, Cardinal RN, Robbins TW (2004) Contrasting roles of basolateral amygdala and orbitofrontal cortex in impulsive choice. J Neurosci Off J Soc Neurosci 24:4718–4722.

Zangemeister L, Grabenhorst F, Schultz W (2016) Neural Basis for Economic Saving Strategies in Human Amygdala-Prefrontal Reward Circuits. Curr Biol 26:3004–3013.

Zhang W, Schneider DM, Belova MA, Morrison SE, Paton JJ, Salzman CD (2013) Functional Circuits and Anatomical Distribution of Response Properties in the Primate Amygdala. J Neurosci 33:722–733.

Zuo Y, Wang X, Cui C, Luo F, Yu P, Wang X (2011) Cocaine-induced Impulsive Choices Are Accompanied by Impaired Delay-dependent Anticipatory Activity in Basolateral Amygdala. J Cogn Neurosci 24:196–211.

